# The Shape of the Bacterial Ribosome Exit Tunnel Affects Cotranslational Protein Folding

**DOI:** 10.1101/274191

**Authors:** Renuka Kudva, Pengfei Tian, Fatima Pardo Avila, Marta Carroni, Robert B. Best, Harris D. Bernstein, Gunnar von Heijne

## Abstract

The *E.coli* ribosome exit tunnel can accommodate small folded proteins, while larger ones fold outside. It remains unclear, however, to what extent the geometry of the tunnel influences protein folding. Here, using *E. coli* ribosomes with deletions in loops in proteins uL23 and uL24 that protrude into the tunnel, we investigate how tunnel geometry determines where proteins of different sizes fold. We find that a 29-residue zinc-finger domain normally folding close to the uL23 loop folds deeper in the tunnel in uL23 Δloop ribosomes, while two ~100-residue protein normally folding close to the uL24 loop near the tunnel exit port fold at deeper locations in uL24 Δloop ribosomes, in good agreement with results obtained by coarse-grained molecular dynamics simulations. This supports the idea that cotranslational folding commences once a protein domain reaches a location in the exit tunnel where there is sufficient space to house the folded structure.

## Introduction

A large fraction of cellular proteins likely start to fold cotranslationally in the ~100 Å long exit tunnel in the ribosomal large subunit (1–3), before they emerge into the cytosolic environment. In *E. coli* ribosomes, portions of the 23S rRNA and a few universally conserved proteins line the exit tunnel, Fig. 1A. The tunnel proteins uL4, uL22, and uL23 consist of globular domains that are buried within the rRNA, and β-hairpin loops that protrude into the tunnel (4). These loops help stabilize the tertiary structure of 23S rRNA (5) and contribute towards the unique geometry of the tunnel (2, 6, 7). uL24 and uL29 are located near the end of the tunnel, and a hairpin loop in uL24 forms a finger-like structure that partially obstructs the tunnel exit port.

**Figure 1.**
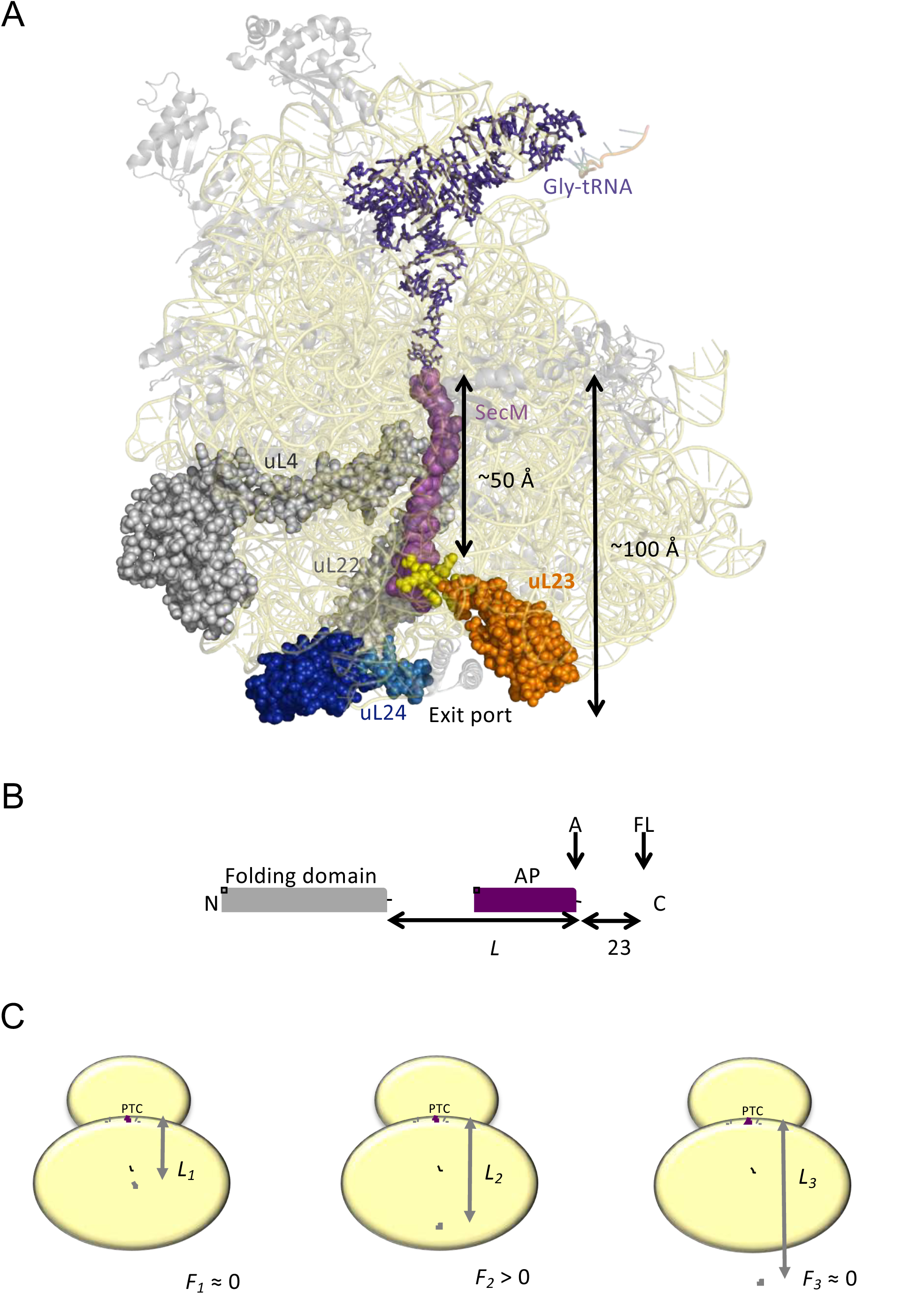
(A) Front view of the 50S subunit of the *E. coli* ribosome adapted from PDB 3JBU (46), with tunnel proteins uL4 and uL22 indicated in gray. The globular domain of uL23 is indicated in orange with the β-hairpin loop depicted in yellow. uL24 is shown in dark blue, with the loop at the tunnel exit shown in light blue. The exit tunnel, outlined by a stalled SecM nascent chain (purple), is ~100 Å in length. (B) The arrest-peptide assay (13). The domain to be studied is placed *L* residues upstream of the critical proline at the C-terminal end of the 17-residue long arrest peptide (AP) from the *E. coli* SecM protein. A 23-residue long stretch of the *E. coli* LepB protein is attached downstream of the AP, allowing us to separate the arrested (A) and full-length (FL) products by SDS-PAGE after translation. Constructs are translated in the PURExpress^®^ *in vitro* translation system supplemented with WT, uL23 Δloop, or uL24 Δloop high-salt washed ribosomes for 20 min. The relative amounts of arrested and full-length protein were estimated by quantification of SDS-PAGE gels, and the fraction of full-length protein was calculated as *f_FL_* = *I_FL_*/(*I_A_* + *I_FL_*) where *I_A_* and *I_FL_* are the intensities of the bands corresponding to the A and FL products. (c) *f_FL_* is a proxy for the force *F* that cotranslational folding of a protein domain exerts on the AP. At short linker lengths, both *F* and *f_FL_* ≈ 0 because the domain is unable to fold due to lack of space in the exit tunnel. At intermediate linker lengths, *F* and *f_FL_* > 0 because the domain pulls on the nascent chain as it folds. At longer linker lengths, *F* and *f_FL_* ≈ 0 because the domain is already folded when the ribosome reaches the end of the AP.

Inspired by observation that protein domains fold in different parts of the exit tunnel depending on their molecular weight (8–12), we now ask what role the geometry of the exit tunnel plays in determining where these domains fold. To explore this question, we employ the same arrest peptide-based approach (and coarse-grained MD simulations) used in our previous studies of cotranslational protein folding (13, 14), but with ribosomes that carry deletions in either the uL23 or the uL24 hairpin loop. Our findings provide strong evidence that the tunnel geometry determines where in the tunnel a protein starts to fold.

## Results and Discussion

### The folding assay

Our experimental set-up, Fig. 1B, exploits the ability of the SecM translational arrest peptide (AP) (15) to act as a force sensor (16–18), making it possible to detect the folding of protein domains in the exit tunnel (13, 18). In brief, the domain to be studied is cloned, via a linker, to the AP, *L* residues away from its C-terminal proline. The AP is followed by a C-terminal tail, to ensure that arrested (*A*) nascent chains can be cleanly separated from full-length (*FL*) chains by SDS-PAGE. Constructs with different *L* are translated in the PURE *in vitro* translation system (19), and the fraction full-length protein (*f_FL_*) is determined for each *L*.

For linkers that, when stretched, are long enough to allow the protein to reach a part of the exit tunnel where it can fold, force will be exerted on the AP by the folding protein, reducing stalling and increasing *f_FL_.* (20), Fig. 1C. A plot of *f_FL_* vs. *L* thus shows where in the exit tunnel a protein starts to fold and at which linker length folding no longer causes increased tension in the nascent chain. A number of earlier studies have provided strong support for the notion that the dominant peak in a *f_FL_* profile corresponds to folding into the native state (as opposed to, *e.g.*, non-specific compaction of the nascent chain), at least for small, single-domain proteins (11, 13, 14, 20–22).

### uL23 Δloop and uL24 Δloop ribosomes

The *E. coli* strains HDB143 (uL23 Δloop; uL23 residues 65-74 deleted) and HDB144 (uL24 Δloop; uL24 residues 43-57 deleted) have previously been shown to be viable (23), as is a strain where uL23 has been replaced by a homologue from spinach chloroplast ribosomes that also lacks the β-hairpin loop (24, 25). These strains were used to purify high-salt-washed ribosomes that were used to translate proteins in the commercially available PURExpress^®^ Δ-Ribosome kit. Analysis of the purified ribosomes by SDS-PAGE and Western blotting demonstrated the expected size differences compared to wildtype for the uL23 Δloop and uL24 Δloop proteins, Figure 1- figure supplement 1.

### Cryo-EM structure of uL23 Δloop ribosomes

The loop deleted in the uL24 Δloop ribosomes does not interact with neighboring parts of the ribosome, Fig. 2A, and hence its removal would not be expected to alter the structure of other parts of the exit tunnel. In contrast, the loop deleted in uL23 Δloop ribosomes is located deep in the exit tunnel, Fig. 2B, ~ 40-50 Å from the exit and it is not clear *a priori* whether its removal may cause rearrangements in other tunnel components. For this reason, we determined a cryoEM structure of the uL23 Δloop 70S ribosome at an average resolution of 3.3 Å, Fig. 2C, Figure 2- figure supplement 1, and found that the shape of the tunnel remains unchanged in the uL23 Δloop ribosome when compared with wildtype (WT) *E. coli* ribosomes, except for an increase in volume resulting from the absence of the uL23 loop Fig. 2C-D. We estimated this increase using the POVME algorithm (26, 27). Compared to WT *E. coli* ribosomes, the tunnel volume increases by 2,064 Å^3^ in uL23 Δloop ribosomes, see Video 1, about 1/3 of the size of ADR1a (5,880 Å^3^) calculated by the same method.

**Figure 2.**
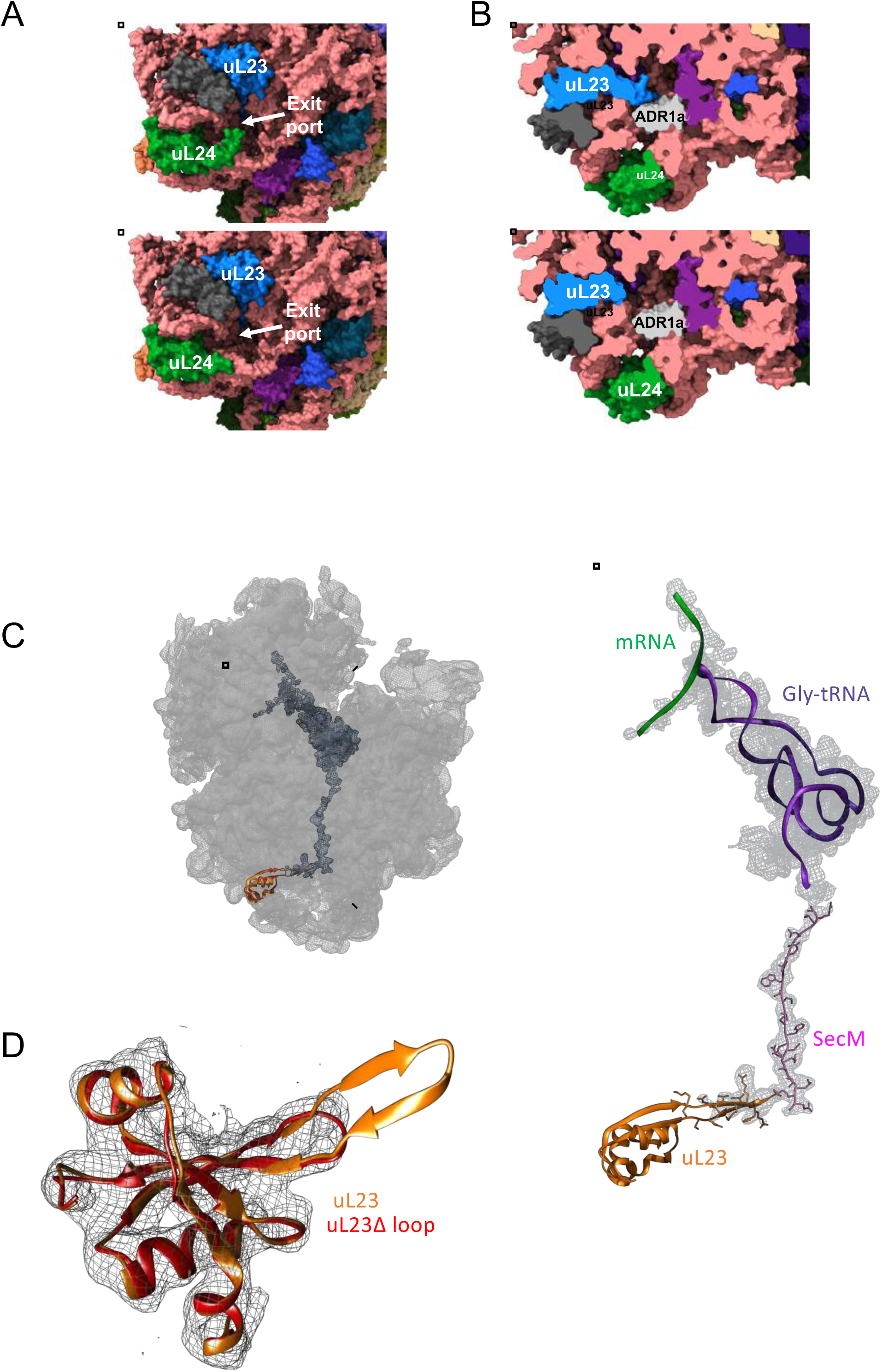
Structural consequences of removing the hairpin loops in uL24 and uL23 modeled after PDB 3JBU of the SecM stalled ribosome. (A) In wildtype ribosomes, the loop in uL24 partially obstructs the exit tunnel (top panel). Its removal in uL24 Δloop ribosomes creates a wide opening into the tunnel (bottom panel). (B) In wildtype ribosomes, the loop in uL23 extends into the exit tunnel (top panel). Its removal in uL23 Δloop ribosomes creates an open space around the area where the ADR1a domain is known to fold (13). The ADR1a structure is from PDB 5A7U. (C) Cryo-EM structure of the uL23 Δloop 70 S ribosome (EMD-4319), fitted to PDB 3JBU (that includes a Gly-tRNA and a 26-residue long arrested SecM AP) to locate uL23 (orange) and the exit tunnel. The enlarged region shows a difference map (in mesh) obtained by subtracting the cryo-EM map of the uL23 Δloop 70 S ribosome from a map generated from 3JBU in Chimera. The difference map shows that the only difference in volume between the two maps is the tRNA (in pink), the SecM AP (in magenta), and the loop deleted from uL23. (D) Extracted cryo-EM density (in mesh) for uL23 in the uL23 Δloop ribosome EMD-4319. Wildtype uL23 (orange) and a *de novo*-built model for the mutant uL23 Δloop protein (PDB 6FU8; red) are shown in ribbon representation.

### ADR1a folds deeper in the exit tunnel in uL23 Δloop but not in uL24 Δloop ribosomes

ADR1a constructs of different linker lengths (*L*) were translated in the PURExpress^®^ Δ-Ribosome kit supplemented with purified WT, uL23 Δloop, or uL24 Δloop ribosomes, either in the presence of 50 μM Zn^2+^ (to promote folding of ADR1a) or in the presence of 50 μM of the zinc-specific chelating agent TPEN (to prevent folding of ADR1a; TPEN is required to remove residual amounts of Zn^2+^ from the PURE lysate) (Figure 3A, Figure 3- figure supplements 1-4). Translation rates in PURE are ~10-fold slower than *in vivo* (28), but since the proteins studied here fold on micro-to-millisecond time scales, i.e., considerably faster than the *in vivo* translation rate, it is safe to assume that the folding reaction has time to equilibrate between each translation step both *in vivo* and in the PURE system.

**Figure 3.**
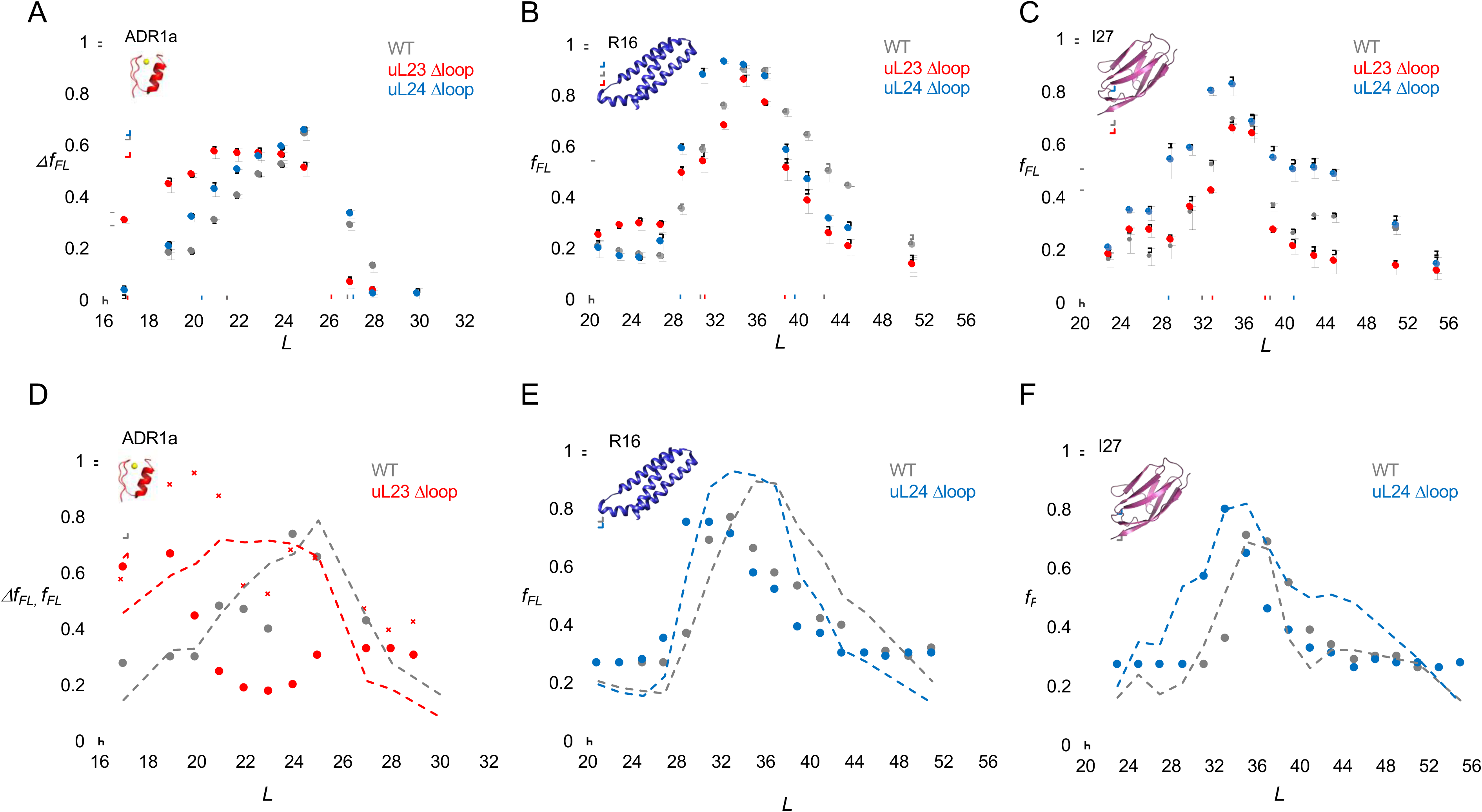
(A) Δ*f_FL_* profiles (Δ*f_FL_* = *f_FL_*(50 μM Zn^2+^) –*f_FL_*(50 μM TPEN)) for ADR1a constructs translated in the PURE system supplemented with in-house purified WT (gray), uL23 Δloop (red), and uL24 Δloop (blue) *E. coli* ribosomes. (B) *f_FL_* profiles for spectrin R16 constructs translated in the PURE system supplemented with in-house purified WT (gray), uL23 Δloop (red), and uL24 Δloop (blue) *E. coli* ribosomes. (C) *f_FL_* profiles for titin I27 constructs translated in the PURE system supplemented with in-house purified WT (gray), uL23 Δloop (red), and uL24 Δloop (blue) *E. coli* ribosomes. Error bars in panels a-c show SEM values calculated from at least 3 independent experiments. Dashed lines indicate *L_onset_* and *L_end_* values, c.f., Table 1. *f_FL_* profiles for non-folding mutants of R16 and I27 are found in (14, 20). (D) Simulated *f_FL_* profiles (full lines) for ADR1a, spectrin R16, and titin I27 obtained with WT (gray), uL23 Δloop (red), and uL24 Δloop (blue) ribosomes. The corresponding experimental *f_FL_* profiles from panels a-c are shown as dashed lines. The simulated ADR1a *f_FL_* profile marked by X’s was obtained with a uL23 Δloop(70-72) ribosome model. Simulated *f_FL_* profiles for ADR1a with uL24 Δloop ribosomes, and for R16 and I27 with uL23 Δloop ribosomes, are essentially identical to the corresponding profiles obtained with WT ribosomes, and are shown in Figure 3-figure supplement 10.

**Table 1.** *L_onset_*, *L_max_*, and *L_end_* values calculated from the *f_FL_* profiles in Fig. 3.

|  | ADR1a |  |  | R16 |  |  | I27 |  |  |
| --- | --- | --- | --- | --- | --- | --- | --- | --- | --- |
| | WT | uL23<br>Δloop | uL24<br>Δloop | WT | uL23<br>Δloop | uL24<br>Δloop | WT | uL23<br>Δloop | uL24<br>Δloop |
| $L_{onset}$ | 21 | 17 | 20 | 31 | 31 | 29 | 32 | 33 | 28 |
| $L_{max}$ | 25 | 22 | 25 | 35 | 35 | 33 | 35 | 35 | 35 |
| $L_{end}$ | 27 | 26 | 27 | 42 | 39 | 40 | 38 | 38 | 41 |

Similar to previous results (13), we saw efficient stalling when ADR1a was translated in the presence of TPEN at *L* ≥ 19 residues, Figure 3- figure supplement 8. Further, there is a slight but significant increase in *f_FL_* at *L* = 17 residues in the presence of TPEN (hence not related to folding); this has been observed before (13) and we hypothesize that it is due to a weakening in the arrest potency of SecM by the ADR1a residues that abut the AP in this construct (see Figure 3- figure supplement 9 for sequences). To correct for this effect, we calculated Δ*f_FL_* = *f_FL_*(Zn^2+^) – *f_FL_*(TPEN), Fig. 3A. In the presence of Zn^2+^, the Δ*f_FL_* profiles for WT and uL24 Δloop ribosomes are very similar: Δ*f_FL_* starts to increase around *L* = 20-21 residues and peaks at *L* = 25 residues (grey and blue curves). In contrast, for the uL23 Δloop ribosomes, Δ*f_FL_* starts to increase already at *L* = 17 residues and peaks at *L* = 21-24 residues (red curve). To quantify these differences, for each *f_FL_* curve we calculated the linker lengths characterizing the onset and end of the peak (*L_onset_* and *L_end_*; defined as the *L*-values for which the curve has half-maximal height, as indicated in Fig. 3A), as well as the *L*-value corresponding to the peak of the curve (*L_max_*), Table 1.

A previous cryo-EM study demonstrated that the 29-residue ADR1a domain folds deep inside the ribosome exit tunnel in a location where it is in contact with the uL23 loop (13), Fig. 2B. The additional space available in uL23 Δloop ribosomes makes it possible for ADR1a to start to fold at 3-4 residues shorter linker lengths (*L_onset_*). Assuming an extended conformation of the linker segment (~3Å per residue), ADR1a folds ~9-12 Å deeper in the exit tunnel in uL23 Δloop ribosomes than in WT ribosomes.

### Spectrin and titin domains fold deeper in the exit tunnel in uL24 Δloop ribosomes

The 109-residue α-spectrin R16 domain has been shown to fold cotranslationally at *L* ≈ 35 residues, in close proximity to uL24 in the exit port region (14). As seen in Fig. 3B and Table 1, with both WT and uL23 Δloop ribosomes, R16 has *L_onset_* = 31 residues and *L_max_* = 35 residues (gray and red curves). For the uL24 Δloop ribosomes however, *L_onset_* = 29 residues and *L_max_* = 33 residues (Table 1), suggesting that that spectrin R16 folds ~6-7 Å deeper in the exit tunnel when the uL24 loop does not obstruct the tunnel exit port.

Similar results were obtained for the 89-residue titin I27 domain, Fig. 3C. Previous studies have shown that the I27 domain folds at linker lengths *L* = 35-39 residues and that it folds in about the same location as does spectrin R16, in close proximity to the uL24 loop (20). The *f_FL_* profile is not affected by the uL23 loop deletion, but folding commences at ~4 residues shorter linker lengths in uL24 Δloop ribosomes, similar to R16 (Table 1).

### Coarse-grained molecular dynamics simulations

In order to provide a more detailed structural framework for interpreting the *f_FL_* profile results, we performed coarse-grained molecular dynamics simulations of the cotranslational folding of ADR1a, spectrin R16, and titin I27 in WT, uL23 Δloop, and uL24 Δloop ribosomes, using a recently described model that allows us to calculate *f_FL_* profiles from the simulations (20). The essence of the method is that the simulations are used to determine folded and unfolded populations at each linker length, and the forces associated with them. Combining this information with the experimentally determined force-dependent escape rate of the AP from the ribosome (18) in a kinetic model allows *f_FL_* to be calculated. Simulated (full lines) and experimental (dashed lines) *f_FL_* profiles are shown in Fig. 3D-F, and detailed simulation results, together with representative snapshots from the simulations of the folded domains at *L* ≈ *L_onset_*, are shown in Figure 3- figure supplement 10. For all three proteins, the *L_onset_* values are well reproduced by the simulations, both in WT and Δloop ribosomes. *L_max_* values are also well captured by the simulations for ADR1a (WT ribosomes) and I27 (both WT and uL24 Δloop ribosomes), but are shifted to somewhat lower values in the R16 simulations.

The simulated ADR1a *f_FL_* profile for uL23 Δloop ribosomes, while showing an early onset of folding in agreement with the experimental profile, has a much smaller *L_max_* value. We also performed a simulation using a ribosome model with a smaller deletion in the uL23 loop (residues 70-72; red curve marked by X’ s); in this case, the peak in the simulated profile extends between *L_onset_* and *L_end_* values that are more similar to the experimental profile for uL23 Δloop ribosomes. The shape of the *f_FL_* profile for ADR1a is clearly highly sensitive to fine structural details of the exit tunnel and therefore somewhat difficult to reproduce by coarse-grained simulations.

In summary, both the experimental and simulation results are consistent with the idea that proteins start to fold as soon as they reach a part of the exit tunnel that is large enough to hold the folded protein. Judging from the *f_FL_* profiles, the 29-residue ADR1a domain folds approximately ~9-12 Å deeper in the exit tunnel in uL23 Δloop ribosomes than in WT and uL24 Δloop ribosomes, while the 89- and 109-residue titin and spectrin domains fold ~6-10 Å deeper inside the tunnel in uL24 Δloop ribosomes than in WT and uL23 Δloop ribosomes; the corresponding values estimated from the simulations are ~6Å for ADR1a and ~13-15 Å for I27 and R16 (Figure 3- figure supplement 10 panel B).

Both the uL23 and uL24 loops thus serve to reduce the space available for folding, but in different parts of the exit tunnel. The uL24 loop is particularly interesting in this regard. In bacterial ribosomes, it partially blocks the tunnel exit port, closing off what would otherwise be a wide, funnel-like opening, Fig. 2a, and thereby prevents domains of M_w_ ≥ 10 kDa from folding inside the exit tunnel. It is conserved (in length, if not in sequence) in bacterial ribosomes, Figure 1- figure supplement 2, suggesting that this will be the case not only for *E. coli* ribosomes but for bacterial ribosomes in general. Eukaryotic ribosome tunnels have different geometries owing to expansion segments in their rRNA as well as an increased number of proteins and a wider exit port (29, 30); uL24 is among the most divergent proteins compared to bacteria (31). We therefore expect the precise relation between the onset of folding and protein M_w_ to be somewhat different in eukaryotic ribosomes, as also suggested by a recent study (32).

## Acknowledgements

This work was supported by grants from the Knut and Alice Wallenberg Foundation, the Swedish Cancer Foundation, and the Swedish Research Council to GvH, and PT, RBB and HDB were supported by the Intramural Research program of the National Institute of Diabetes and Digestive and Kidney Diseases of the NIH. FPA was supported by NIH grant R35GM122543. The cryo-EM data were collected at the Swedish National Cryo-EM Facility funded by the Knut and Alice Wallenberg Foundation, the Family Erling Persson Foundation and the Science for Life Laboratory. This work utilized the computational resources of the NIH HPC Biowulf cluster (http://hpc.nih.gov).

## Materials and Methods

Key Resources

| Reagent type (species) or resource | Designation | Source or reference | Identifiers | Additional information |
| --- | --- | --- | --- | --- |
| strain, strain background (*Escherichia coli*) | HDB140, HDB143, HDB144, Strain background MC4100 | 10.1111/j.1365-2958.2010.07325.x | CGSC#: 6152 | Strains used to isolate high-salt washed ribosomes in this study. |
| antibody | uL23, uL24 | 10.1111/j.1365-2958.2010.07325.x | NA | 1:8000 dilution used (incubated for one hour). Secondary antibody: Mouse (1:20,000 dilution incubated for one hour). Nitrocellulose membrane blocked with 5% Milk in TBS-T for an hour. |
| peptide, recombinant protein | uL23 $\Delta$ loop, uL24 $\Delta$ loop | 10.1111/j.1365-2958.2010.07325.x | UniProtKB-P0ADZ0 (rplW)<br>UniProtKB-P60624 (rplX) | Referred to as HDB 143 and HDB 144 in original paper. Refers to genes rplWΔ65-74 and rplXΔ43-57 respectively. |
| commercial assay or kit | GeneJET Plasmid miniprep kit | Thermo Fisher Scientific<br>RRID:SCR_008452 | Cat no. K0502 | Used to purify plasmids |
| commercial assay or kit | PURExpress<sup>®</sup> $\Delta$ Ribosome kit | New England Biolabs | Cat no. E3313S | Kit was supplemented with ribosomes purified in the lab. |
|  |  |  |  | Translation carried out at 37°C for 20 min. |
| software, algorithm | EasyQuant | doi: 10.1038/nsmb.2376 | NA | Used to quantify relative fraction full-length of translated protein from SDS-PAGE |
| software, algorithm | cryoSPARC version v2 | Structura Biotechnology Inc | NA | Used for ab-initio reconstruction of uL23Δloop ribosomes. The following operations were carried out as part of the cryoSPARC workflow: 2D classification, Ab initio reconstruction, Homogeneous refinement, Sharpening and map flipping to correct for handedness, local resolution. |
| software, algorithm | UCSF ChimeraX |  | SCR_015872 | Used to make Figure 2A |
| software, algorithm | UCSF Chimera v. 1.12 | J Comput Chem. 2004 Oct;25(13):1605-12. | SCR_004097 | Used to visualise the cryoEM map, fit PDB models 3JBU, 4YBB to check for differences in maps. Used to make figures. |
| software, algorithm | Jalview v 2.10.4 | doi: 10.1093/bioinformatics/btp033 | SCR_006459 | Use for generating multiple sequence alignments of uL23 and uL24 in the supplementary figures. |
| chemical | Potassium acetate | Sigma-Aldrich (SCR_008988) | Cat no. P1190 | Source of potassium ions to stabilise ribosomes |
| chemical | Magnesium acetate | Sigma-Aldrich (SCR_008988) | Cat no. M5661 | Source of Magnesium ions to stabilise ribosomes |
| chemical | cOmplete protease | Sigma-Aldrich | Cat no. 04693116001 | Used as a protease inhibitor during cell |
|  | inhibitor cocktail | (SCR_008988) |  | lysis to obtain ribosomes |
| chemical | N,N,N',N'-Tetrakis(2-pyridylmethyl)ethylenediamine | Sigma Aldrich (SCR_008988) | Cat no. P4413 | Used to chelate Zn for the -Zn reactions in the ADR1 plot. |
| chemical | threo-1,4-Dimercapto-2,3-butanediol DL-Dithiothreitol | Sigma Aldrich (SCR_008988) | Cat no. DTT-RO | Reducing agent added to buffers during ribosome purification and as a reductant for SDS-PAGE |
| chemical | Tris Base | Sigma-Aldrich (SCR_008988) | Cat no. T1503 | Buffering agent during ribosome preparation |
| chemical | <sup>35</sup>S Methionine | Perkin-Elmer | Cat no. NEG009T001MC | <sup>35</sup>S Methionine is incorporated into the protein during in vitro translation and aids detection by phosphorimaging. |

### Enzymes and Chemicals

The PURExpress^®^ Δ Ribosome kit was purchased from New England Biolabs (Cat no. E3313S). The components used to prepare Lysogeny Broth (LB Medium) for ribosome isolation were obtained from BD Biosciences and all other chemicals used were sourced from Merck Sigma Aldrich. (^35^S) Methionine was purchased from Perkin Elmer. Bis-Tris gels and plasmid isolation kits were obtained from Thermo Scientific.

### Plasmids

All ADR1, spectrin and titin constructs fused to the *E. coli* SecM AP via a variable linker were expressed from the pET19b vector, as described previously (13, 14, 20). The spectrin constructs used in this study lacked the soluble domain of LepB at the N-terminus.

### Strains and antisera

Strains HDB140 (referred to as WT), HDB143 (referred to as uL23 Δloop) and HDB144 (referred to as uL24 Δloop), as well as rabbit polyclonal antisera against uL23 and uL24, are described in (23).

### Isolation of ribosomes

Ribosomes were purified from the strains HDB140, HDB143, and HDB144. The strains were cultured in Lysogeny broth (LB) to an A_600_ of 1.0 at 37°C and chilled on ice for 15 min before they were harvested by centrifugation at 4000 g for 10 min. The cell pellet was washed twice with Buffer A at pH 7.5 (10 mM Tris-OAc, 14 mM Mg(OAc)_2_, 60 mM KOAc, 1 mM DTT, 0.1% Complete Protease Inhibitor) and lysed using the Emulsifex (Avestin) at a pressure of 8000 psi. The cell lysate was loaded on a sucrose cushion at pH 7.5 (50 mM Tris-OAc, 1 M KOAc, 15 mM Mg(OAc)_2_,1.44 M Sucrose, 1 mM DTT, 0.1% Complete Protease Inhibitor) and centrifuged at 80,000xg in a Ti70 rotor (Beckman-Coulter) for 17 hours. The obtained ribosomal pellet was resuspended in Buffer B at pH 7.5 (50 mM Tris-OAc, 50 mM KOAc, 5 mM Mg(OAc)_2_, 1 mM DTT), flask frozen in liquid nitrogen and stored at -80°C. This suspension of ribosomes is presumed to consist of a pool of non-translating 30S, 50S and 70S particles due to the concentration of Mg^2+^ in the buffer they are in. Each batch of ribosomes that was prepared was tested for optimal translation by titrating different volumes in the PURExpress^®^ Δ-Ribosome kit.

### In vitro transcription and translation

The generated constructs were translated for 20 min. in the PURExpress^®^ Δ-Ribosome kit supplemented with high-salt-washed ribosomes isolated from HDB140, HDB143 (uL23 Δloop), or HDB144 (uL24 Δloop). Plasmid DNA of each construct (300 ng) was used as a template for polypeptide synthesis, and translation was carried out in the presence of (^35^S) Methionine at 37°C for 20 min and shaking at 500 r.p.m. For ADR1a constructs, the translation reactions also included either 50 μM ZnCl_2_ or 50 μM of the Zn^2+^ chelator TPEN. Translation was stopped by treating the sample with a final concentration of 5% trichloroacetic acid (TCA) and incubated on ice for 30 min. The TCA precipitated samples were subsequently centrifuged at 20,000 g for 10 min in a tabletop centrifuge (Eppendorf) and the pellet obtained was solubilized in sample buffer, supplemented with RNaseA (400 μg/ml), and incubated at 37°C for 15 minutes. The samples were resolved on 12% Bis-Tris gels (Thermo Scientific) in MOPS buffer for ADR1 and MES buffer for Spectrin and Titin. Gels were dried and subjected to autoradiography and scanned using the Fujifilm FLA-9000 phosphorimager for visualization of radioactively labeled translated proteins.

### Quantification of radioactively labelled proteins

The protein bands on the gel were quantified using MultiGauge (Fujifilm) from which one-dimensional intensity profiles of each gel lane was extracted. This information was subsequently fit to a Gaussian distribution using EasyQuant (Rickard Hedman, Stockholm University). The sum of the arrested and full-length bands was calculated, and this was used to estimate the fraction full-length protein for each construct.

### Cryo-EM sample preparation and data processing

The uL23 Δloop ribosomes (4 A_260_/ml) diluted in grid buffer (20 mM HEPES-KOH, 50 mM KOAc, 10 mM Mg(OAc)_2_, 125 mM sucrose, 2 mM Trp, 0.03% DDM ) were loaded on Pelco^®^ TEM 400 mesh Cu grids pre-coated with 2 nm thick carbon and frozen using the Vitrobot Mark IV (FEI). Data was collected on the Titan Krios (FEI) microscope operated at 300 keV and equipped with a Falcon II direct electron detector. The camera was set to a nominal magnification of 75,000X, which resulted in a pixel size of 1.09 Å at the sample level and a defocus range of -1 to -3 μm.

The frame dose used was 1.17 e/Å^2^, and 20 frames were aligned using MotionCor2 (33) within the Scipion software suite (34). The micrographs were visually inspected and those within a resolution threshold of 5 Å were selected, yielding 3522 micrographs.

471272 particles were picked using Xmipp manual-pick followed by particle extraction within Scipion and further processing in CryoSPARC (35). Two rounds of 2D classification were done, and particles resembling 30S and 50S subunits alone were discarded after visual inspection of the classes. The remaining 297,363 particles of the 70S ribosome were subjected to *ab initio* reconstruction into 3 classes to further sort out heterogeneity. A single homogeneous class consisting of 132,029 particles was used for final homogeneous refinement that resulted in a final map with an average FSC resolution at 0.143 of 3.28 Å. The obtained map of the 70S ribosome was sharpened and corrected for handedness in CryoSPARC fitted with PDBs 3JBU and 4YBB in Chimera (36). Local resolution and FSC at 0.143 was estimated in cryoSPARC. The electron microscopy map was deposited in the Electron Microscopy Data Bank.

The initial model for uL23 Δloop was built with Coot, and improved by energy minimization in a solvated dodecahedron box of explicit TIP3P waters, neutralized with chloride ions and using the Amber 99SB-ILDN force field (37). The steepest descent minimization method implemented in GROMACS 2016.1 was used (38) (39). Even after minimization, the backbone of the new loop formed after the deletion of residues 65-74 still showed improper geometry and Ramachandran outliers, so we used kinematic sampling (40) to model alternative loop conformations, and then we selected the loop that could fit the electron density and had the best Ramachandran score.

Figures were prepared using MacPymol 1.8.6.2 (Schrödinger LLC), Chimera (36), and ChimeraX (41).

### Calculation of tunnel volume

The volume calculations were performed with POVME 2.0 (1). We used the *E. coli* SecM structure PDB 3JBU as our reference. To determine the inclusion region, we generated a series of overlapping spheres-eight with a 20 Å radius, and one with a 40 Å radius. In order to have a complete coverage of the exit tunnel, the centers of the spheres were chosen to match the coordinates corresponding to alternating Ca atoms of the amino acids of the SecM arrest peptide located within the exit tunnel (for the 20 Å radius spheres the residues use as centers were D11, F13, T15, V17, I19, Q21, Q23, I25, A27, G28 and for the 40 Å radius sphere the residue was E3). Grid Spacing was set to 2.0 Å, and the distance cut-off to 1.09 Å. For all three cases (WT, uL24 Δ loop, uL23 Δ loop), we used the same inclusion region. We also removed the SecM arrest peptide located within the exit tunnel. For uL23 Δloop ribosomes residues 65-75 were removed from uL23, and for uL24 Δloop ribosomes residues 42-57 were removed from uL24 (numbering based on PDB 3JBU) prior to the calculation.

### Kinetic model to calculate fraction full length protein f_FL_(t)

The theoretical force profiles (Fig. 3d-f) for ADR1a, I27, and R16 were calculated based on a kinetic model introduced in our previous study (20). Briefly, the rate, k_e_, of the arrest peptide sequence escape from the peptidyl transfer center with a force (F) exerted by the folding protein can be calculated using the Bell model:

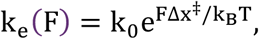

where ΔX^‡^ is the distance from the free energy minimum to the transition state, k_0_ the rupture rate when force equals to zero, k_B_ is Boltzmann’s constant, and T the absolute temperature. In this study, k_0_ and Δx^‡^ are set to be 3.4 × 10^-4^ s^-1^ and 4.5 Å, respectively, based on a previous experimental study (18) in which k_0_ and Δx^‡^ were estimated to be in the range of 0.5 × 10^-4^ to 20 × 10^-4^ s^-1^ and 1-8 Å, respectively.

We assume that the folding and unfolding of the protein is much faster than the escape from the ribosome. Then the time-dependent force profile f_FL_(t) can be obtained approximately by the mean pulling forces exerted when the protein is unfolded, F_u_, or folded, F_f_, and the unfolded and folded populations of P_u_ and P_f_ respectively,

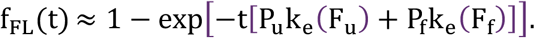

Note that the values of F_u_, F_f_, P_u_, P_f_ are dependent on the linker length *L*, and can be determined by molecular dynamics simulations.

### Molecular dynamics simulations of ribosome-nascent chain complex

A coarse-grained model was employed to simulate folding of ADR1a, I27, and R16 on the ribosome. The ribosome was modelled on the 50S subunit of the *E. coli* ribosome (PDB 3OFR; (42)) Each amino acid in the nascent chain and ribosome was represented by one bead at the position of the α-carbon atom, each RNA residue was modelled by three beads which represent phosphate P, sugar C4′, and base N3 (43). The uL23 Δloop ribosome was modelled by replacing the coordinates of the wildtype uL23 protein with the cryo-EM structure of the uL23 Δloop protein from this study (PDB 6FU8), after being aligned to the wild type protein. The uL24 Δloop ribosome was modelled by replacing the coordinates of the wildtype uL24 protein with the structure of the uL24 Δloop protein built by homology modelling with Modeller (44).

The interactions within the nascent chain were governed by a standard structure-based model (45), which allowed it to reversibly fold to the native state and unfold. Interactions between the protein and ribosome beads were purely repulsive. The pulling force (F) exerted on the arrest peptide by the folding of the protein (ADR1a, R16, or I27) was measured by the extension of the harmonic pseudobond potential between the last and the second last amino acid of the SecM arrest peptide. More details can be found in our previous study (20).

## Figure legends

**Figure 1- figure supplement 1.**
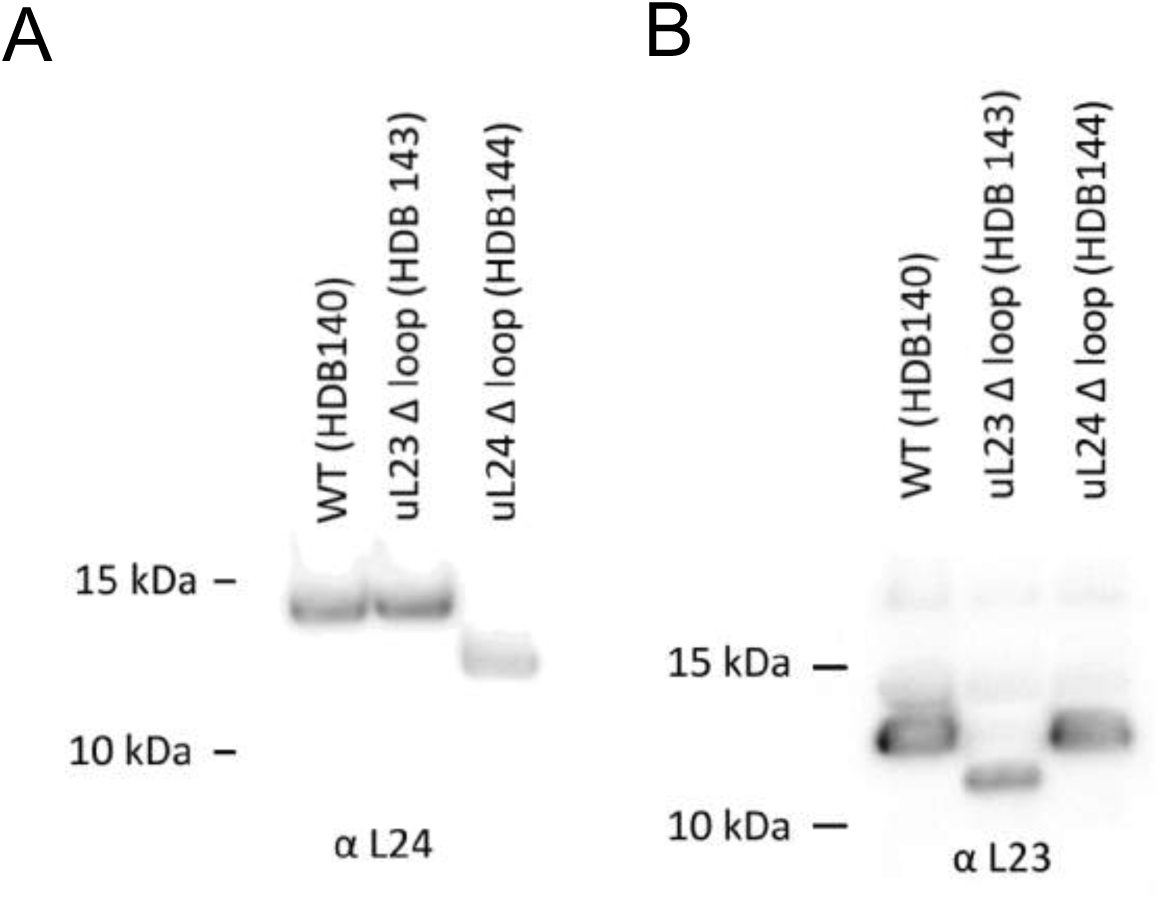
A_260_ = 300 units (6.9 μM) of high-salt washed ribosomes were separated on a 12% Bis-Tris gel and transferred by Western blotting onto a nitrocellulose membrane and detected with antibodies against uL24 (panel A) or uL23 (panel B).

**Figure 1- figure supplement 2.**
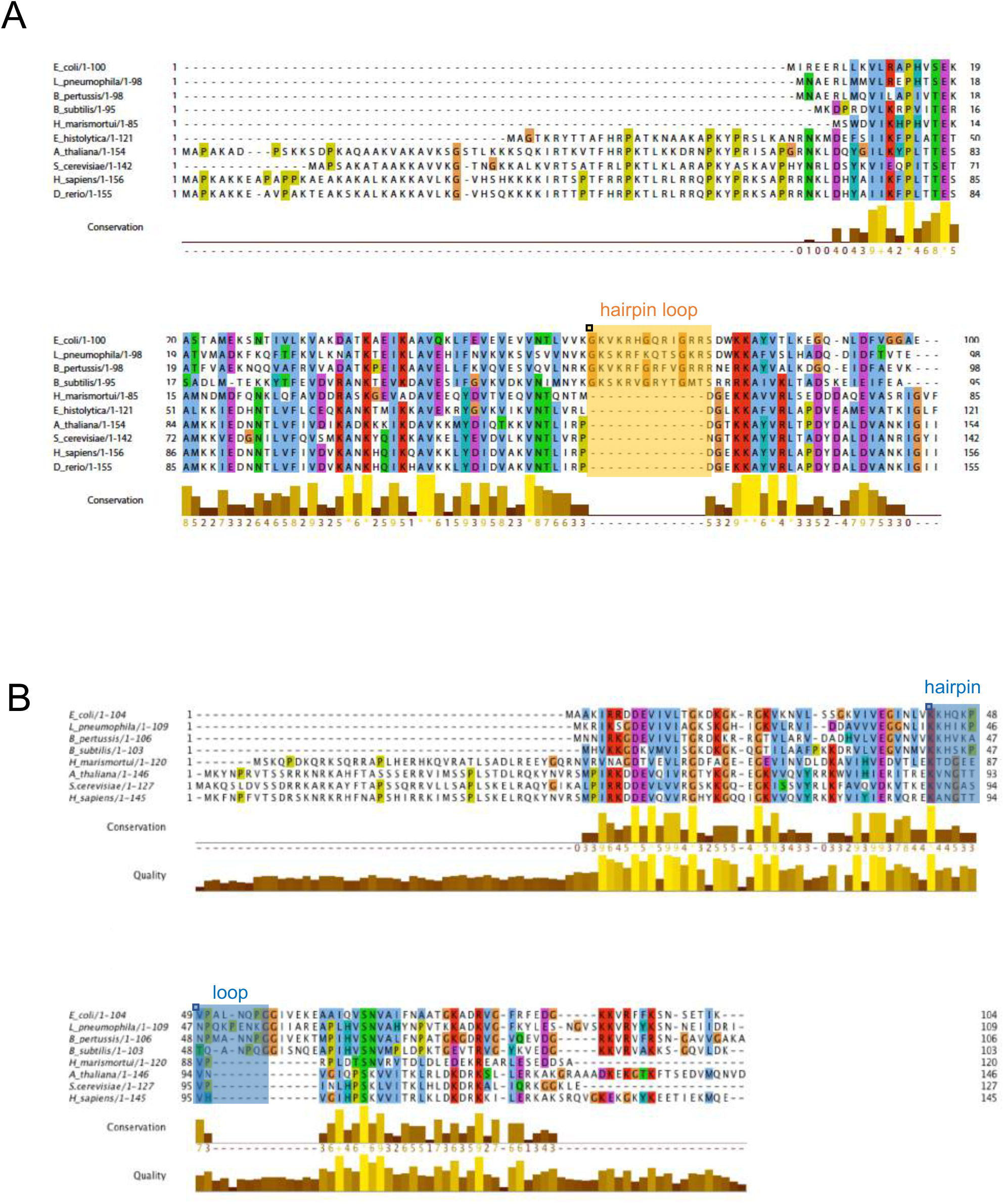
Multiple sequence alignment of uL23 (A) and uL24 (B). The uL23 and uL24 b-hairpin loops, boxed, respectively, in orange and blue, are conserved among Gram-negative bacteria, but are short or absent in archaea and eukaryotes. In eukaryotes and archaea, part of the function of uL23 is proposed to be fulfilled by ribosomal protein eL39.

**Figure 2- figure supplement 1.**
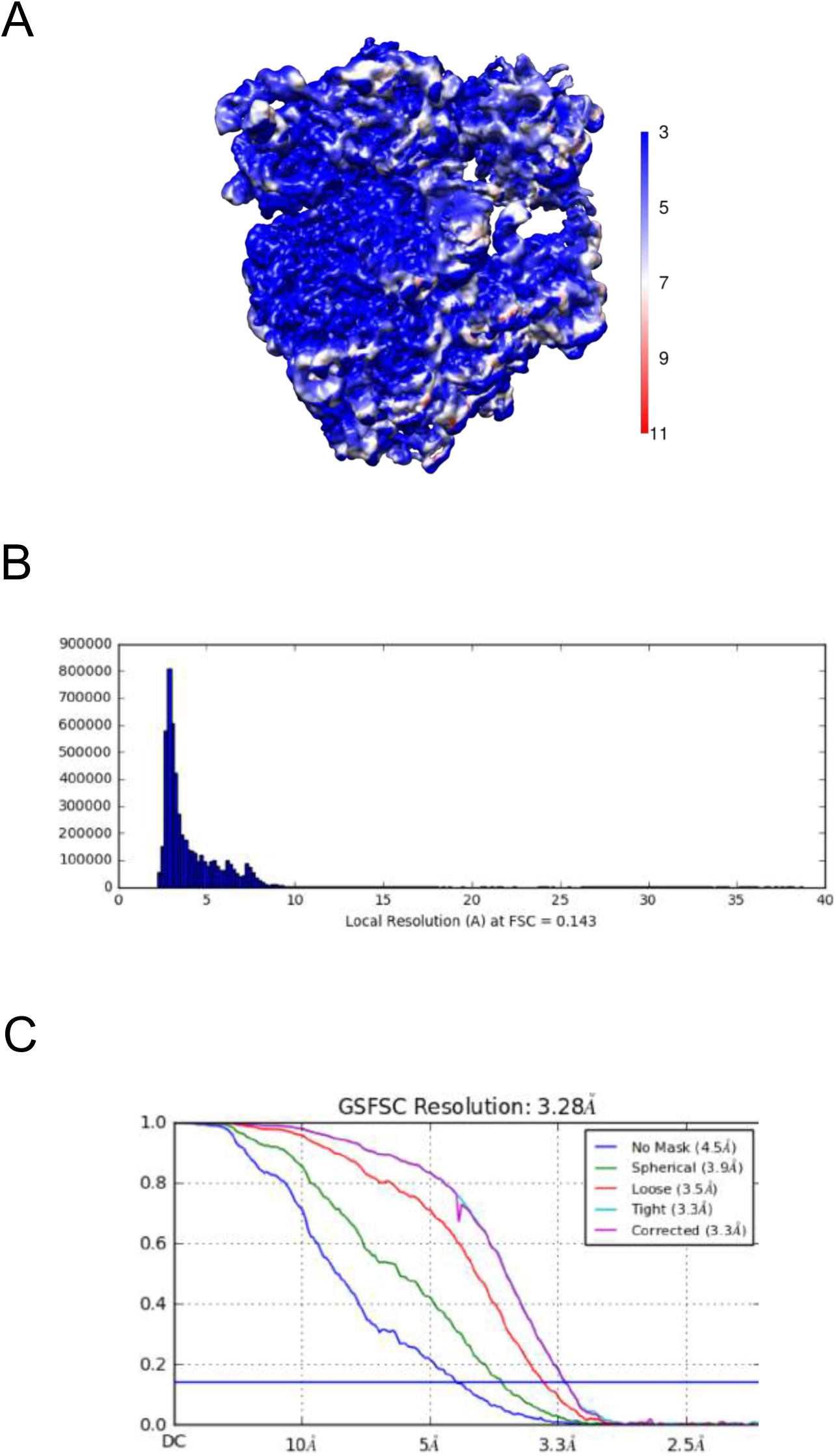
Resolution map of the uL23Δ loop ribosome. (A) Calculation of the local resolution using cryoSPARC. (B) Resolution histogram at FSC 0.143. (C) Fourier-Shell correlation (FSC) curve of the refined map at 0.143 indicating an average resolution of 3.28 Å.

**Figure 3- figure supplement 1-4.**
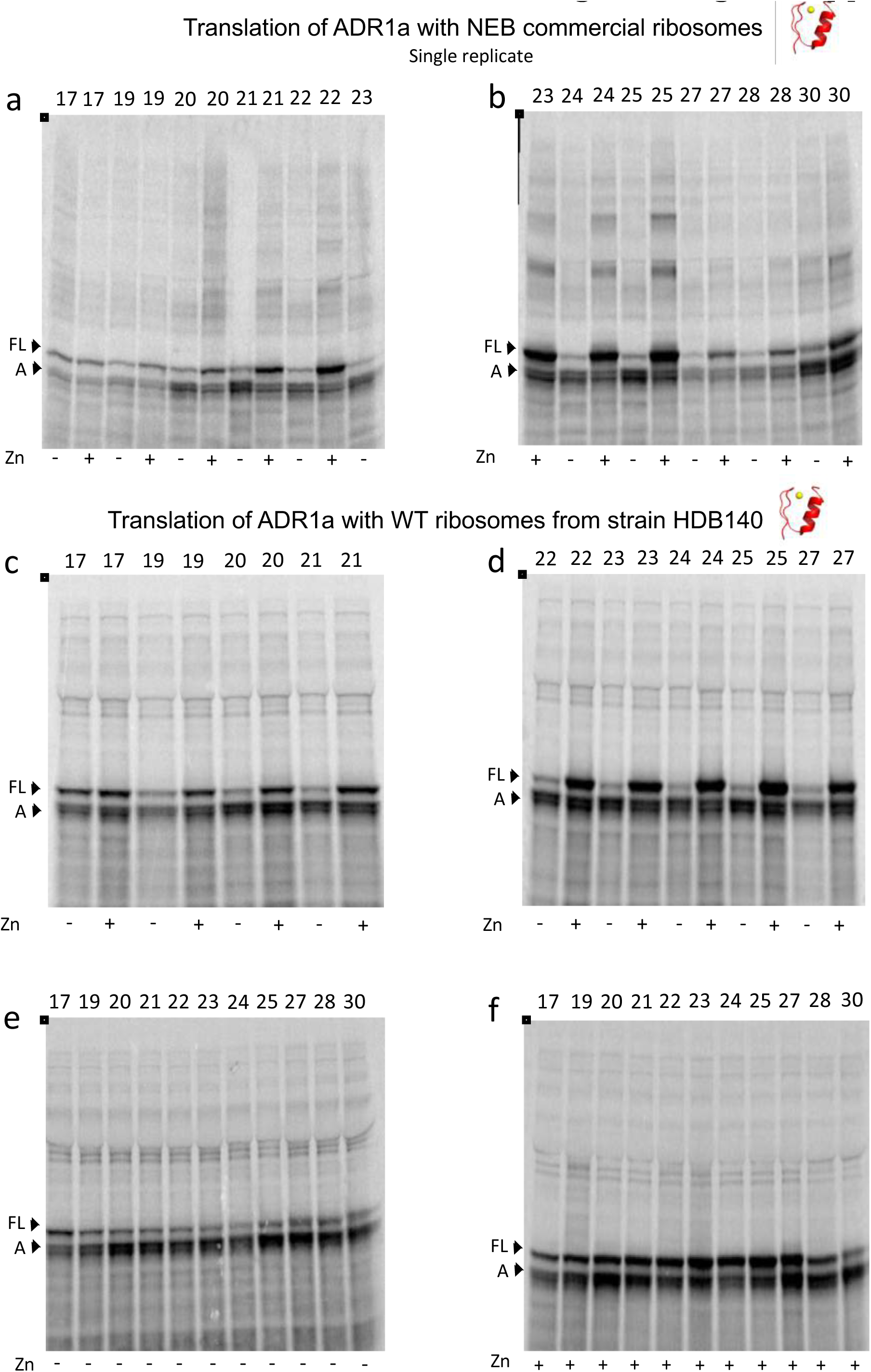
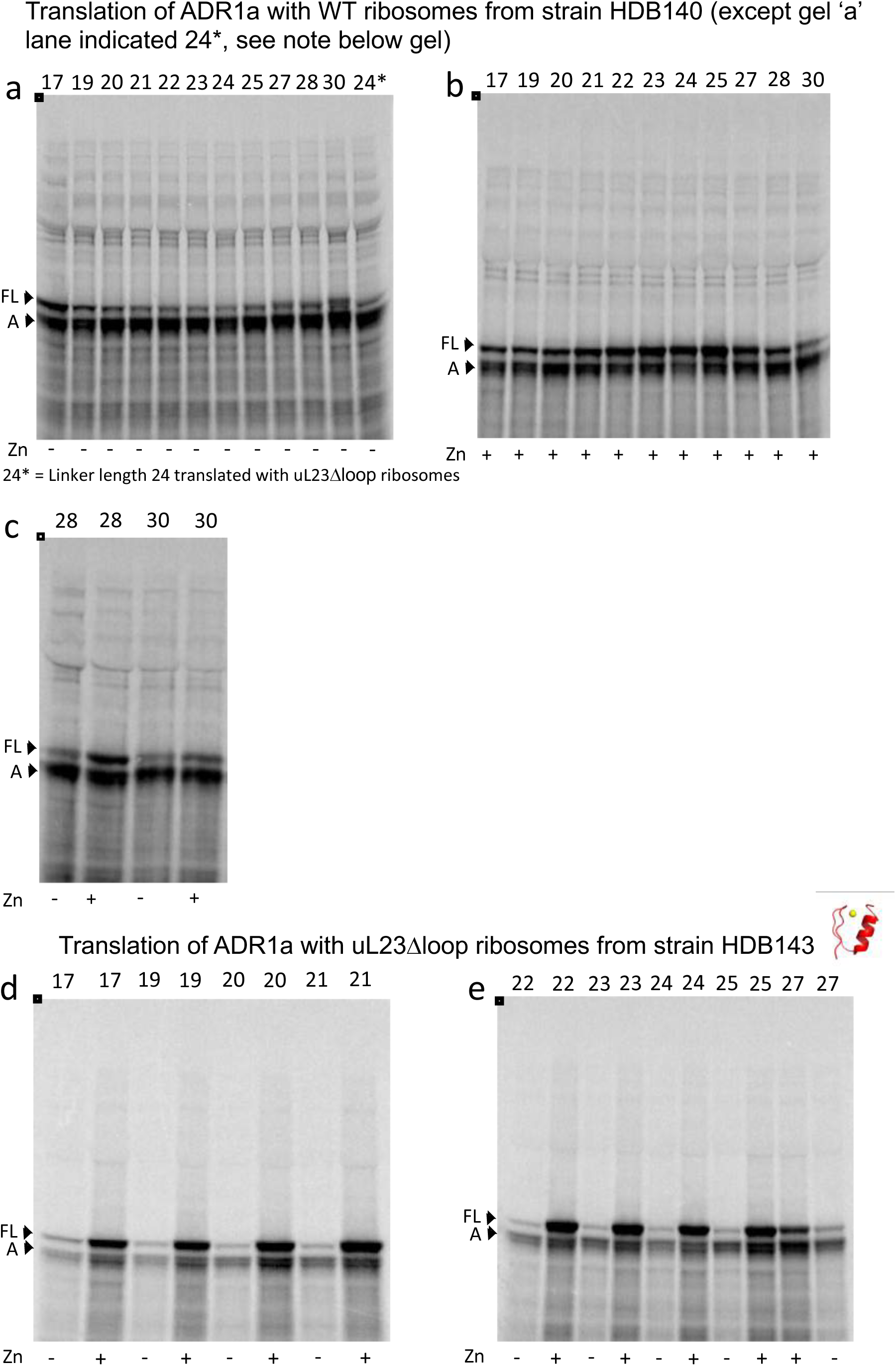
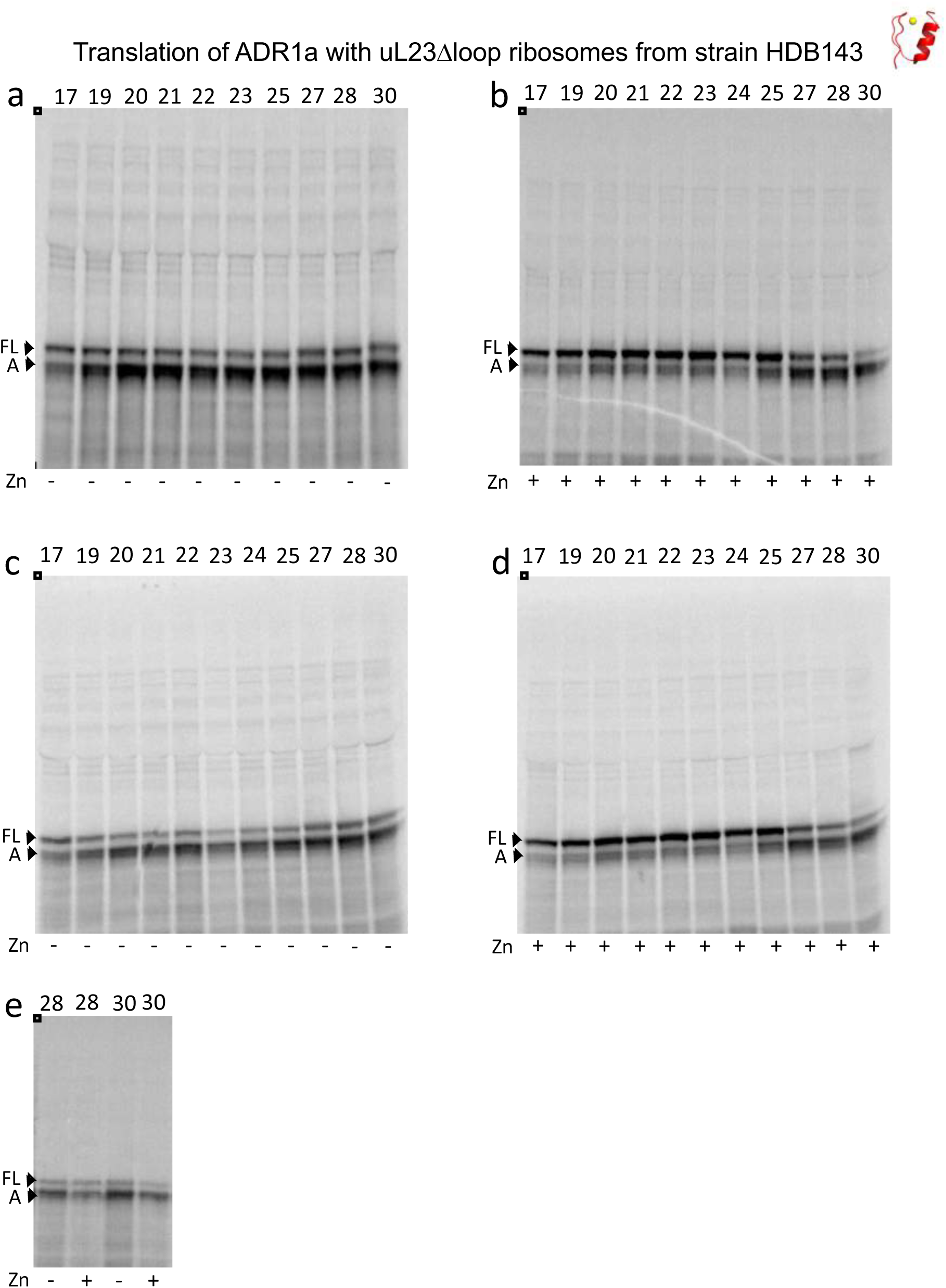
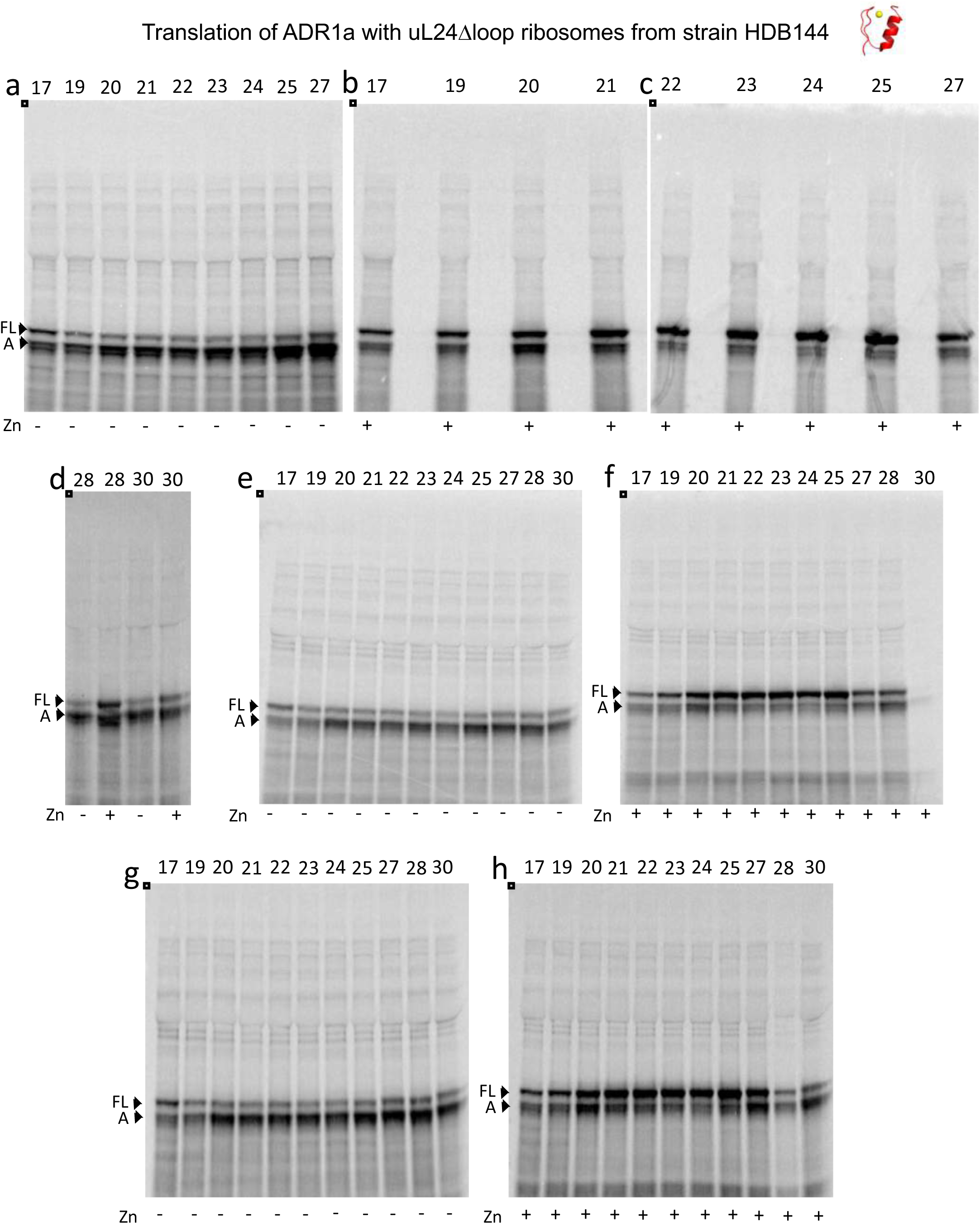
SDS PAGE showing ADR1 constructs translated in PURExpress^®^ Δ-Ribosome kit supplemented with high-salt-washed ribosomes isolated from HDB140, HDB143 (uL23 Δloop), or HDB144 (uL24 Δloop) as indicated. Translations were run on 12% Bis-Tris gels with MOPS running buffer.

**Figure 3- figure supplement 5-6.**
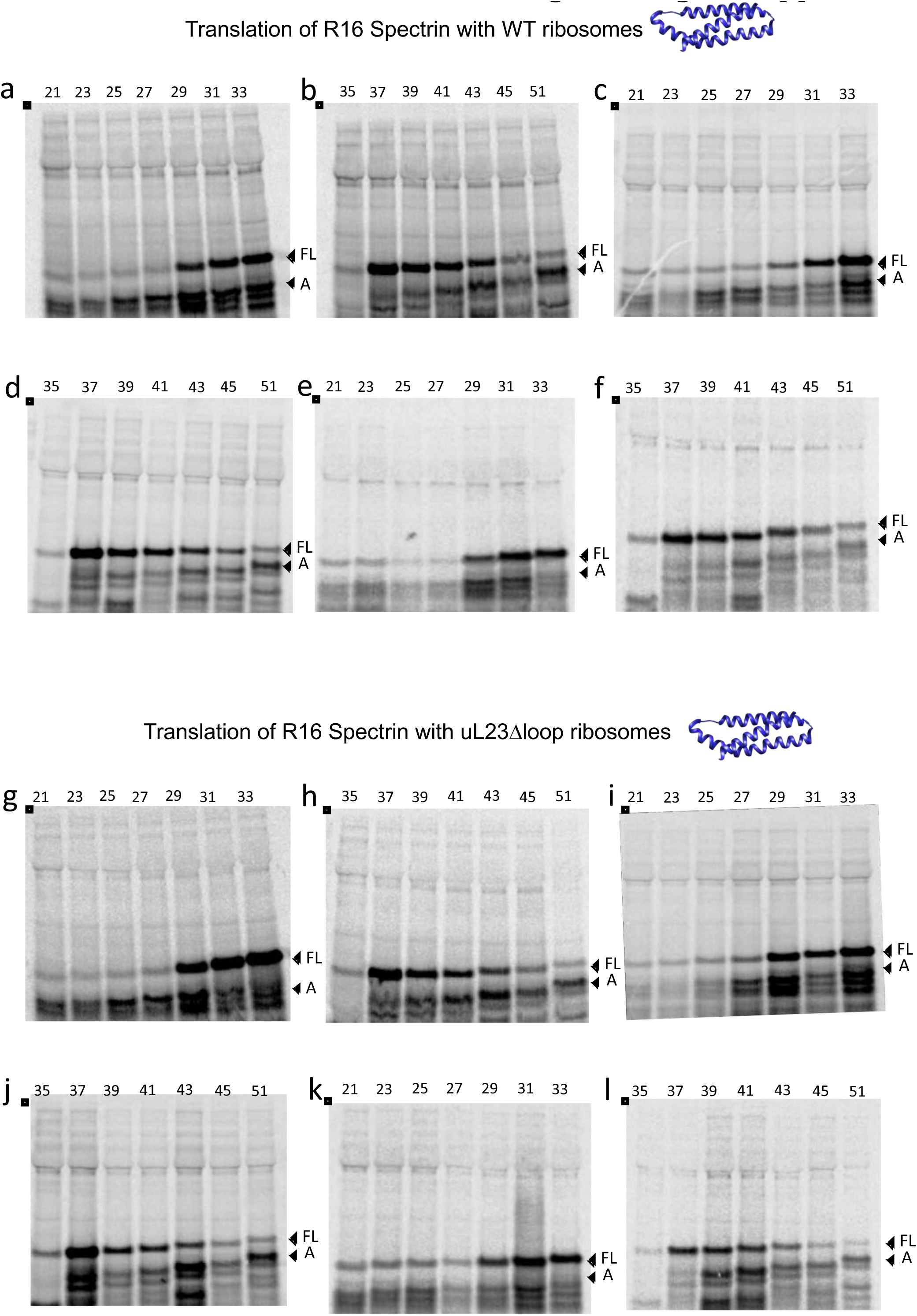
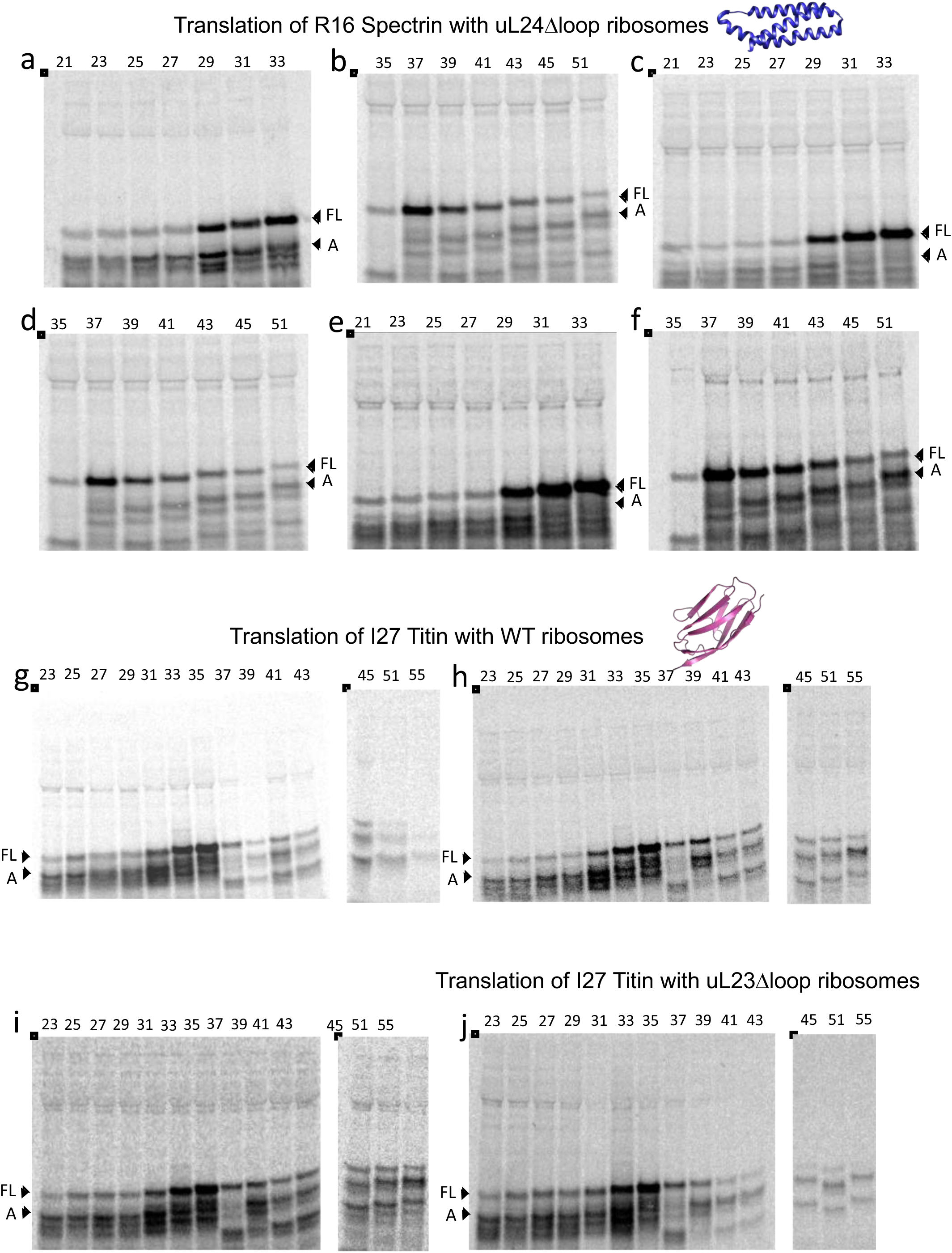
SDS PAGE showing Spectrin R16 constructs translated in PURExpress^®^ Δ-Ribosome kit supplemented with high-salt-washed ribosomes isolated from HDB140, HDB143 (uL23 Δloop), or HDB144 (uL24 Δloop) as indicated. Translations were run on 12% Bis-Tris gels with MES running buffer.

**Figure 3- figure supplement 6-7.**
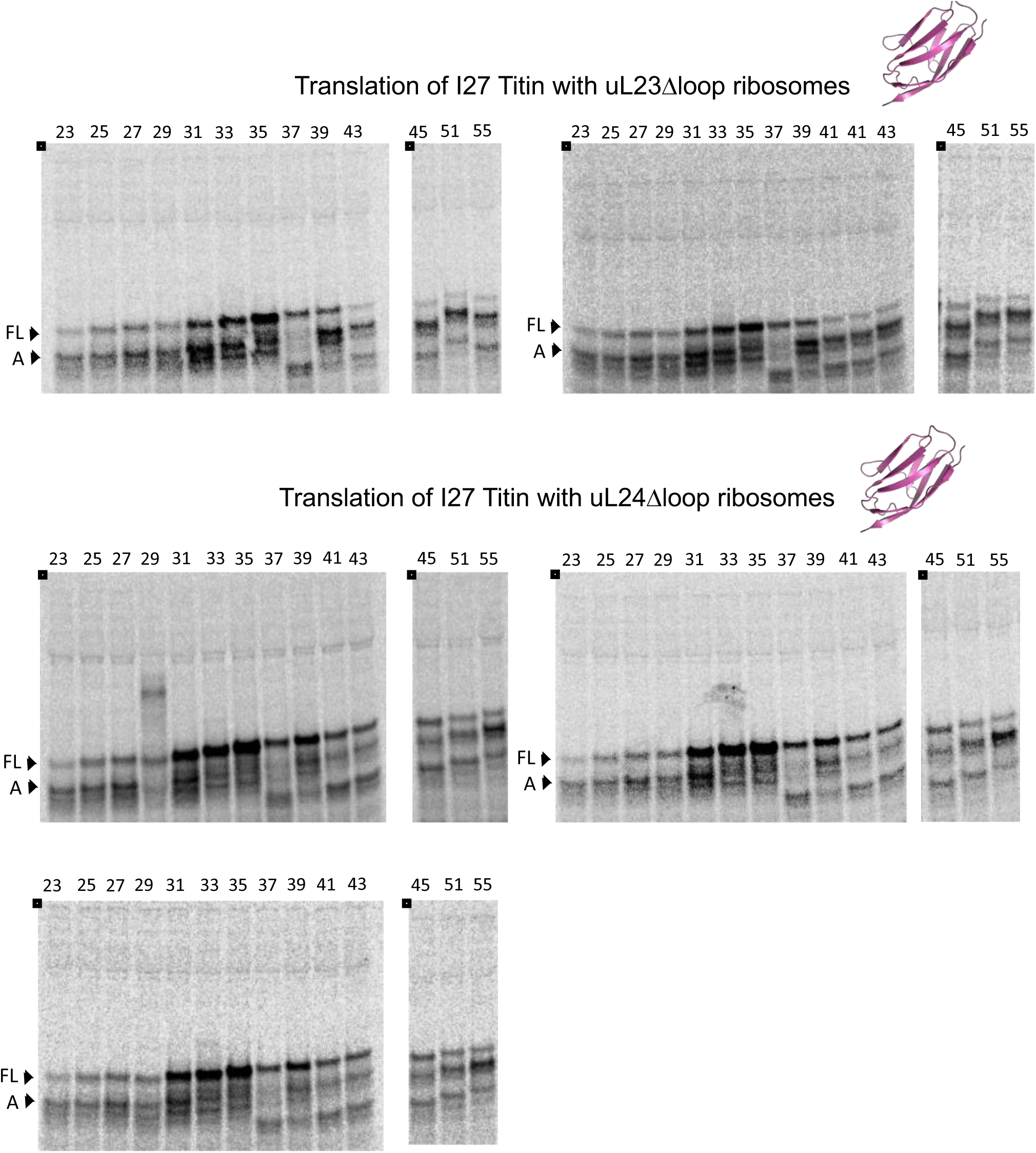
SDS PAGE showing Titin I27 constructs translated in PURExpress^®^ Δ-Ribosome kit supplemented with high-salt-washed ribosomes isolated from HDB140, HDB143 (uL23 Δloop), or HDB144 (uL24 Δloop) as indicated. Translations were run on 12% Bis-Tris gels with MES running buffer.

**Figure 3- figure supplement 8.**
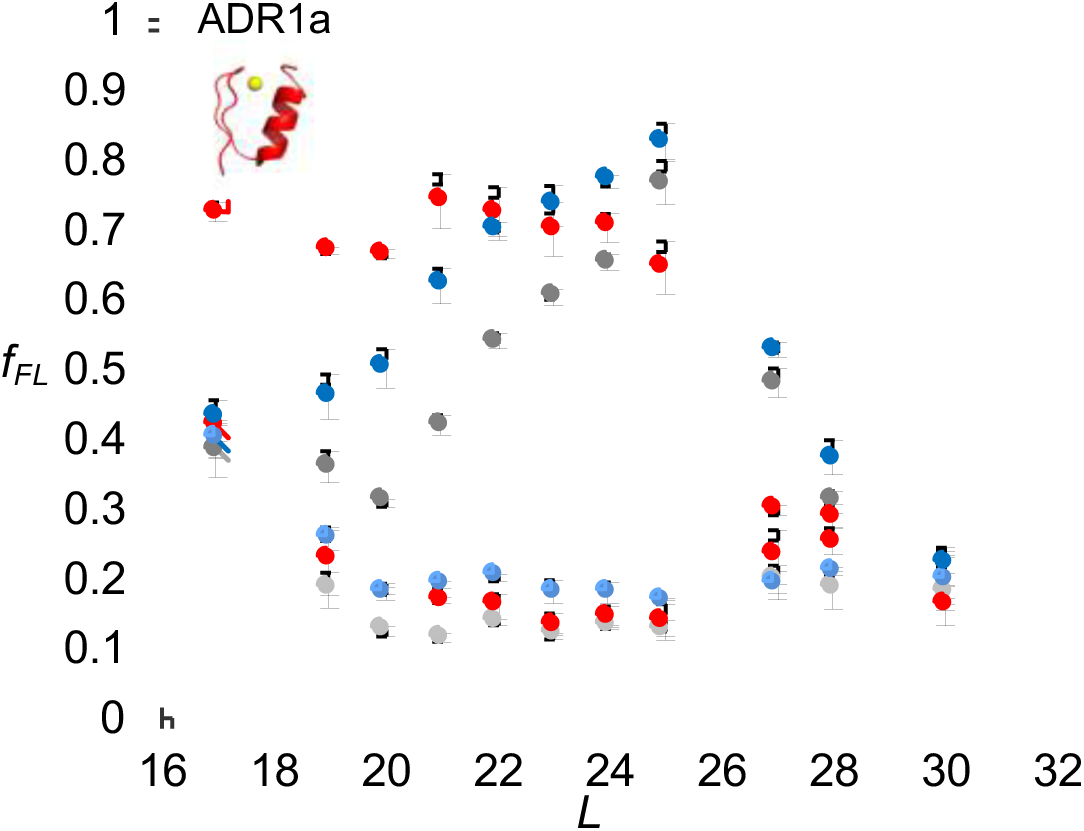
*f_FL_* profiles for ADR1a constructs translated in PURE by WT, uL23 Δloop, and uL24 Δloop ribosomes, either in the presence of Zn^2+^ or of the Zn^2+^ chelator TPEN. Averages and standard errors calculated from three independent translation reactions are shown.

**Figure 3- figure supplement 9.**
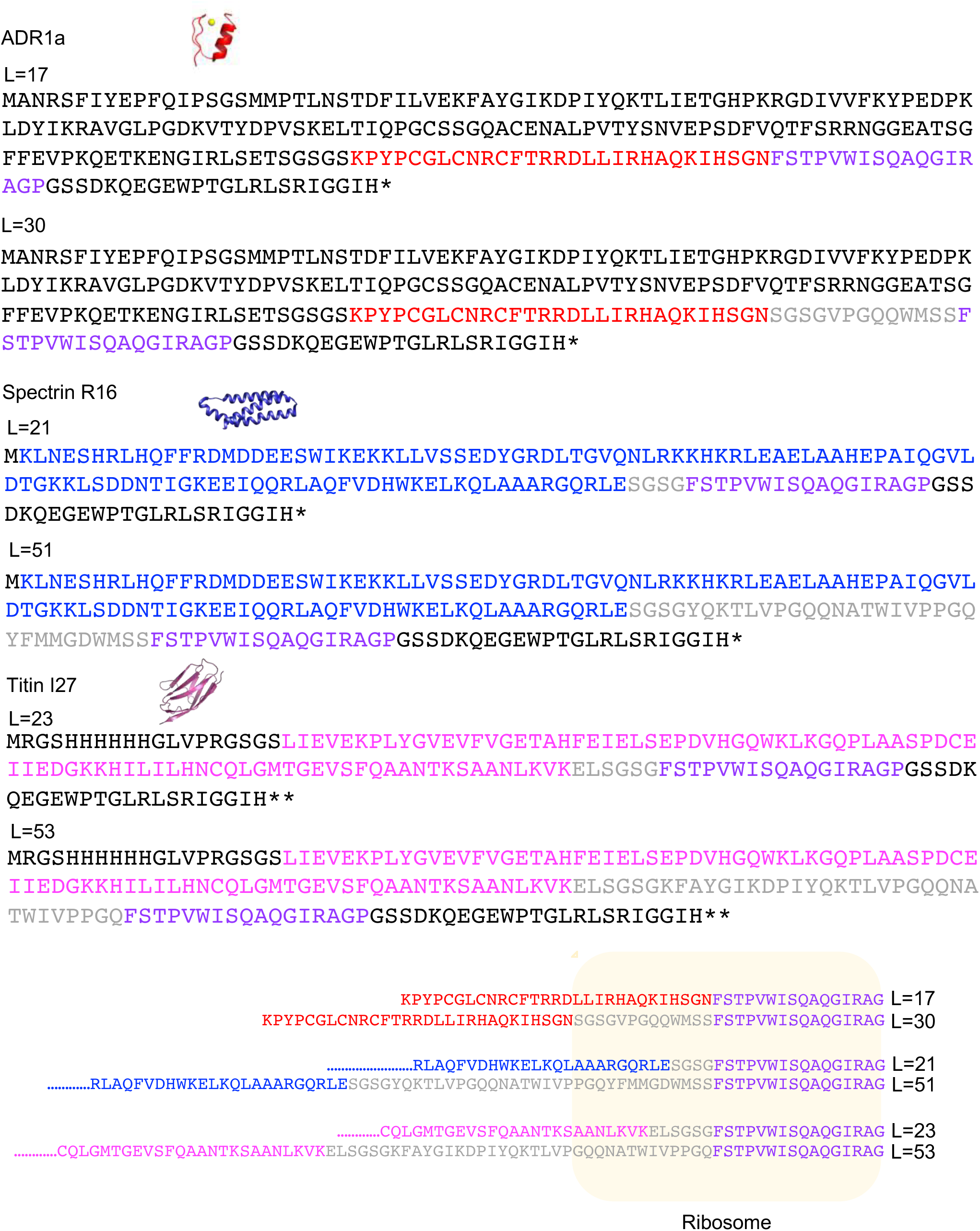
Sequences of the longest and the shortest constructs used for each protein and, a depiction of the location of the sequences the ribosome exit tunnel when the last residue of the AP is in the P-site (lower panel, yellow box). ADR1 is indicated in red, spectrin R16 in blue, titin I27 in pink, and the SecM AP in magenta. The linker between each domain and the AP is in grey. The part of LepB added to the N-terminus of ADR1a is in black.

**Figure 3- figure supplement 10.**
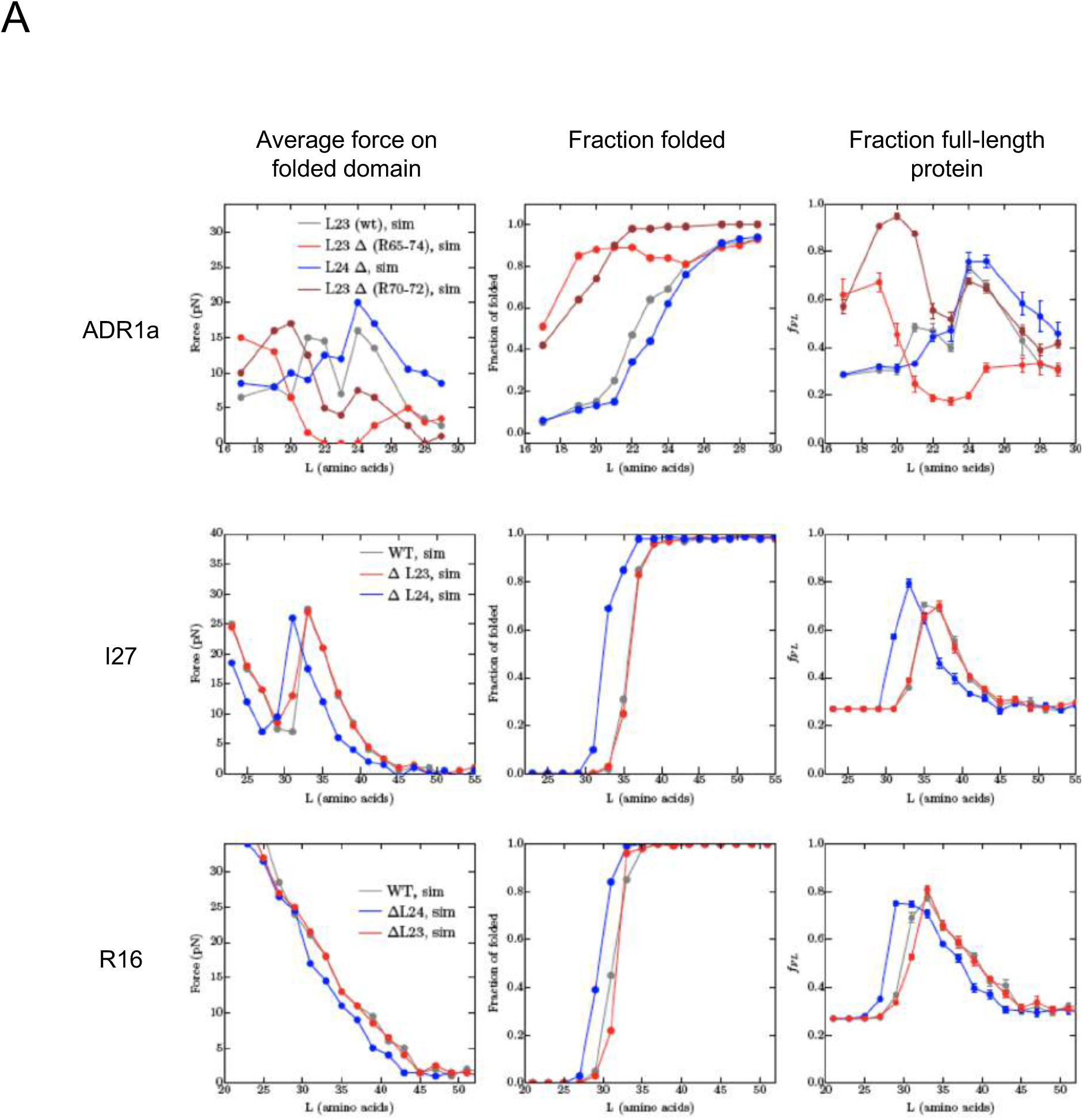
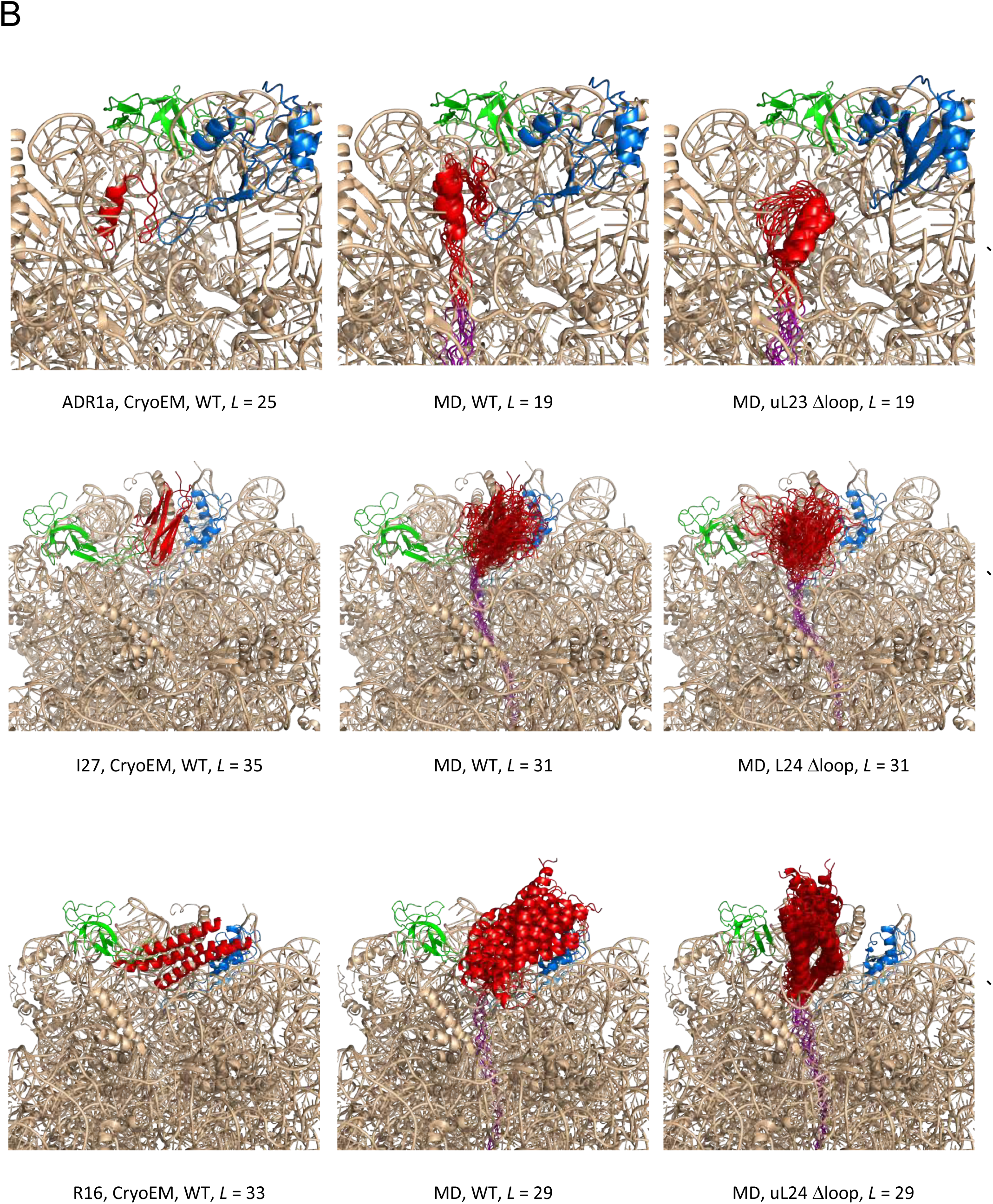
Summary of results from coarse-grained MD simulations. (A) Average forces exerted on the AP by the folded state (first column), fraction folded protein (second column), and *f_FL_* values (last column) for ADR1a, I27, and R16 at different linker lengths *L.* (B) Snapshots of folded ADR1a, I27, and R16 domains in wildtype (WT), uL23 Δloop, and uL24 Δloop ribosomes at *L* values ≈ *L_onset_*. Note that the folded proteins are located at similar depths in the exit tunnel in the cryo-EM structures and the simulations for WT ribosomes (this holds also for ADR1a in uL24 Δloop ribosomes as well as for I27 and R16 in uL23 Δloop ribosomes, c.f. panel A). The C terminus of folded ADR1a is located ~6 Å deeper in the exit tunnel in uL23 Δloop than in WT ribosomes, while folded I27 and R16 are located, respectively, ~15 Å and ~13 Å deeper in the exit tunnel in uL24 Δloop ribosomes (dashed guide lines). Note that the linker is more stretched in the WT ribosome simulations compared to the Δloop ribosomes, consistent with the higher force and lower fraction folded protein seen for the WT ribosome data at the same *L* values (panel a).

*Video 1.* The ribosome exit tunnel (mesh), as calculated for PDB 3JBU, uL23 Δloop and uL24 Δloop ribosomes by POVME. See Fig. 1A for coloring scheme. The b-hairpin loops deleted in uL23 Δloop and uL24 Δloop ribosomes are shown in yellow and light blue, respectively. To facilitate the visualization of the exit tunnel, spheres left outside the exit tunnel after POVME processing were manually removed.

## Abbreviations

AP =: arrest peptide
PTC =: Peptidyl Transferase Centre

